# Medium-chain fatty acids mitigate endoplasmic reticulum stress in yeast cells

**DOI:** 10.1101/2025.09.02.673703

**Authors:** Yuki Ishiwata-Kimata, Yukio Kimata

## Abstract

Upon endoplasmic reticulum (ER) stress, eukaryotic cells commonly trigger cytoprotective transcriptome changes, namely the unfolded protein response (UPR). In yeast *Saccharomyces cerevisiae*, the UPR is mediated by the transcription factor Hac1, which is induced in response to ER stress. Since Hac1 controls hundreds of genes, the biological phenomena that result from UPR are not yet fully understood. Here, we show that cells carrying a mutation to constitutively express Hac1 abundantly contained C10:0 and C12:0 fatty acids, known as medium-chain fatty acids (MCFs). UPR induction by some ER stress stimuli was attenuated by externally supplied MCFs in cells in which fatty acid elongation was genetically or pharmacologically halted. The cell survival assay also indicated the mitigation of ER stress by the MCFs. Moreover, we demonstrated that MCFs leads to the diffusion of a mutant transmembrane protein aggregated in the ER. We propose that, as a biologically beneficial outcome of UPR, MCFs are produced to change the properties of the ER membrane, such as fluidity, in ER-stressed yeast cells.

## INTRODUCTION

Endoplasmic reticulum (ER) is a cellular compartment in which secretory and transmembrane proteins are folded, modified, and assembled before being transported to the Golgi apparatus. Dysfunction of the ER, namely ER stress, is frequently accompanied by the accumulation of unfolded ER client proteins in the ER and provokes a cytoprotective transcriptome change known as the unfolded protein response (UPR). Whereas the UPR is commonly observed in a wide variety of eukaryotic species, its molecular mechanism was initially uncovered through frontier studies using the yeast *Saccharomyces cerevisiae* as a model organism (Ishiwata-Kimata and Kimata, 2023).

Ire1 is an ER-located transmembrane protein that is evolutionarily conserved in eukaryotic species and acts as an ER stress sensor that triggers the UPR. In response to ER stress, Ire1 is self-associated and acts as an endoribonuclease. In *S. cerevisiae* cells, Ire1 promotes the splicing of *HAC1* mRNA to generate the intronless form, hereafter called *HAC1*i mRNA (i stands for “induced”), which is translated into the nuclear transcription factor Hac1. On the other hand, the unspliced form of *HAC1* mRNA, hereafter called *HAC1*u mRNA (u stands for “uninduced”), is deduced to be functionless. Transcriptome analysis of *S. cerevisiae* demonstrated that Hac1 induces or represses hundreds of genes (Travers et al., 2000; Kimata et al., 2006; Ishiwata-Kimata et al., 2025). The most prominent category of UPR targets upregulated by Hac1 are genes encoding ER-localized molecular chaperones and enzymes involved in the modification of ER-client proteins. The UPR also stimulates ER-associated protein degradation (ERAD), through which ER-accumulated unfolded proteins are retrieved to the cytosol and digested by the proteasome (Friedlander et al., 2000; Travers et al., 2000; Hwang and Qi, 2018). These observations explain a role of the UPR in coping with disturbance in protein integrity in the ER. Moreover, Hac1 induces genes related to phospholipid biogenesis and causes ER expansion, which also contributes to efficient protein folding in the ER (Schuck et al., 2009; Nguyen PTM et al., 2022; Niemelä et al., 2024; Ishiwata-Kimata et al., 2025).

However, since Hac1 controls a large number of genes, we speculated that yet unknown outcomes may result from the Hac1-dependent gene regulation in the UPR. Whereas Hac1 increases the cellular abundance of lipid molecules (Nguyen et al., 2022; Ishiwata-Kimata et al., 2025), it remains unclear how the UPR influences lipid composition. Here, we show a considerable abundance of medium-chain fatty acids (MCFs) in strongly UPR-induced *S. cerevisiae* cells. We deduce that this phenomenon is physiologically meaningful because MCFs can mitigate ER stress.

## RESULTS

We previously generated an *S. cerevisiae* mutant, hereafter called *HAC1*i cells, which carried a genomic mutation to remove the intron sequence of the *HAC1* gene (Kimata et al., 2006; Nguyen et al., 2022). Because *HAC1*i cells constitutively express Hac1, leading to unregulated induction of the UPR, biological phenomena induced by the UPR can be easily assessed without exposing cells to external stress stimuli. To investigate the effect of UPR induction on lipid properties in *S. cerevisiae*, here we compared the fatty acid compositions of total lipid samples extracted from *HAC1*i cells and control wild-type cells (Fig. 1). As commonly known, the fatty acid samples were mainly composed of palmitic acid (C16:0), stearic acid (C18:0), palmitoleic acid (C16:1), and oleic acid (C18:1) (Fig. 1). However, we also found that, unlike wild-type cells, *HAC1*i cells contained capric acid (C10:0) and lauric acid (C12:0), known as MCFs, at considerable levels.

**Figure 1.**
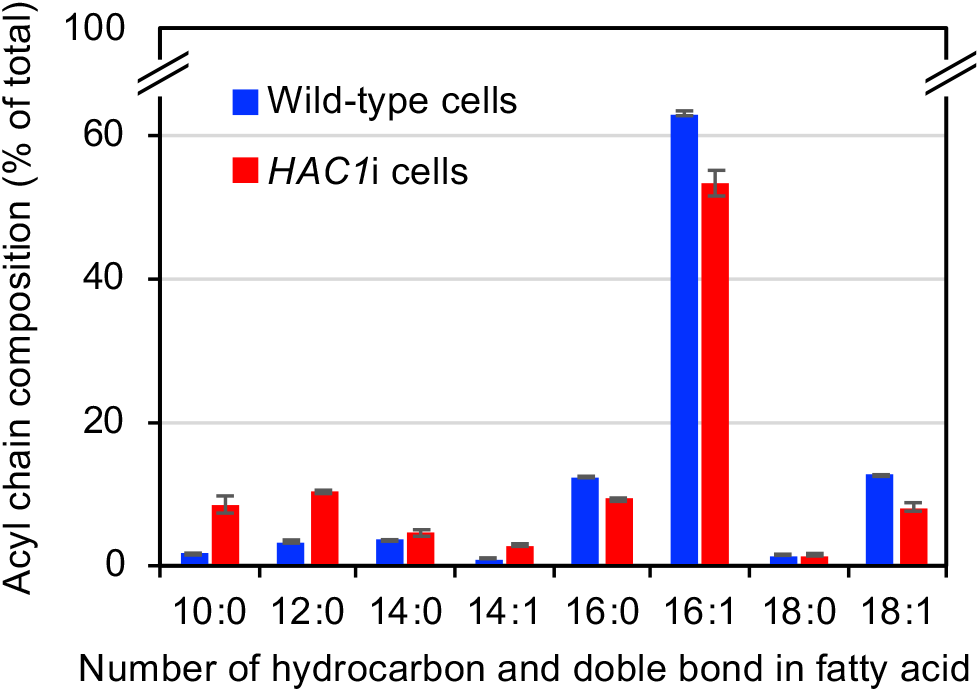
High MCF levels in *HAC1*i cells. Wild-type (BY4742) and *HAC1*i cells were cultured in SC and examined for lipid fatty-acid composition.

We therefore deduced that MCFs are generated as an outcome of the Ire1/Hac1-dependent UPR signaling pathway, and investigated whether this phenomenon contributes to the mitigation of ER stress. To assess cellular functions of MCFs incorporated into lipid molecules, we modified the experimental procedures shown in Otoguro et al. (1981) and Toke and Martin (1996). In *S. cerevisiae* cells, the fatty-acid elongation is mainly performed by the Fas1/Fas2-containing fatty acid synthase complex, which is inactivated by the antifungal antibiotic cerulenin. Thus, extracellularly added MCFs are not converted to longer fatty acids cells treated with cerulenin or carrying the *FAS1* or *FAS2* knockout mutation.

Here, we treated cells with cerulenin in the presence of palmitic acid (16:0), which supports cellular growth when the fatty-acid elongation is blocked, and further added capric acid (C10:0) or lauric acid (C12:0) to the culture media. As conventionally done, ER stress and UPR levels were monitored by *HAC1* mRNA splicing. Fig. 2A and B show that *HAC1* mRNA splicing induced by the N-glycosylation inhibitor tunicamycin or the expression of an ER-accumulating mutant of the multi-membrane-spanning protein Pma1 (Pma1-2308-mCherry; Mai et al., 2019) was drastically suppressed by the extracellularly added MCFs. Although less substantially, MCFs also compromised *HAC1* mRNA splicing caused by inositol depletion, which is known to induce ER stress in *S. cerevisiae* (Fig. 2C). Fig. 2D shows suppression of the tunicamycin-induced *HAC1* mRNA splicing by MCFs in *FAS2* knockout mutant (*fas2Δ*) cells.

**Figure 2.**
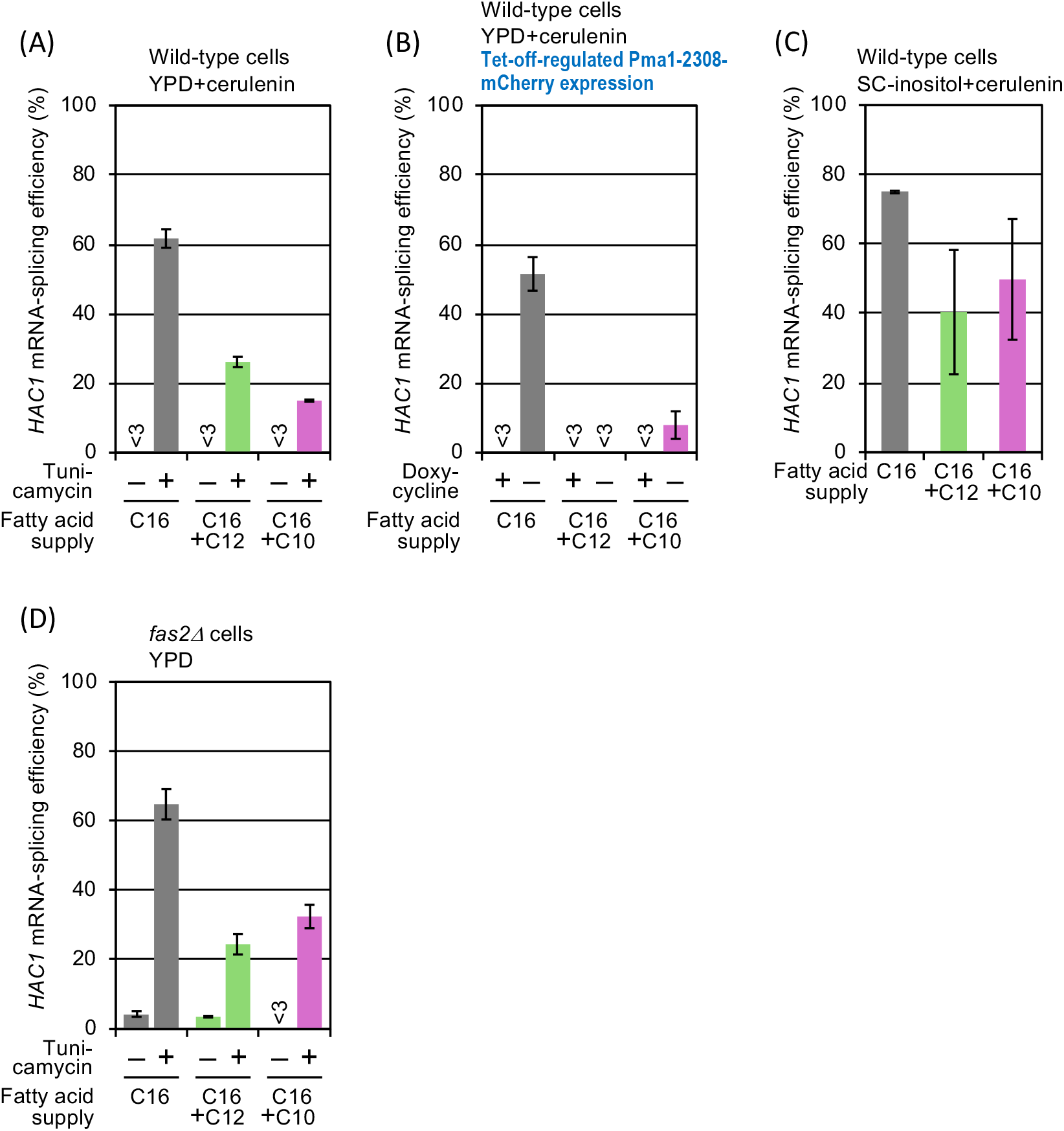
Attenuation of *HAC1* mRNA splicing by MCFs upon various ER stress conditions. (A) Wild-type cells (BY4741) were cultured in fatty acid-supplied YPD containing 5.0 µg/mL cerulenin and were (or were not) treated with 0.6 µg/mL tunicamycin for 1 hr. (B) Wild-type cells (BY4741) carrying pCM189-PMA1-2308-mCherry were cultured in fatty acid-supplied YPD containing 5.0 µg/mL cerulenin in the absence or presence of 3.0 µg/mL doxycycline, which represses the Tet-off promoter-dependent expression of Pma1-2308-mCherry. (C) Wild-type cells (BY4741) were cultured in fatty acid-supplied SC-iniositol for 5hrs. (D) *fas2Δ* cells (BY4741-*fas2Δ*) were cultured in fatty acid-supplied YPD and were (or were not) treated with 0.6 µg/mL tunicamycin for 1 hr. The fatty acids supplied were 500 µg/mL palmitic acid only (C16), 130 µM lauric acid with 500 µg/mL palmitic acid (C16+C12), or 130 µM capric acid with 500 µg/mL palmitic acid (C16+C10). Then, splicing of *HAC1* mRNA was monitored and presented.

Because the UPR is halted, *IRE1* knockout mutant (*ire1Δ*) cells exhibit hypersensitivity to ER stress stimuli including tunicamycin. As shown in Fig. 3A, the viability of tunicamycin-treated *ire1Δ* cells was rescued by the MCF treatment at least partially. On the other hands, MCFs did not seem to affect the viability of tunicamycin-treated *IRE1+* cells (Fig. 3B), although, as aforementioned, *HAC1* mRNA splicing was suppressed by MCFs under this condition (Fig. 2A). This observation suggests that ER stress aggravated by the *ire1Δ* mutation can be alleviated by MCFs.

**Figure 3.**
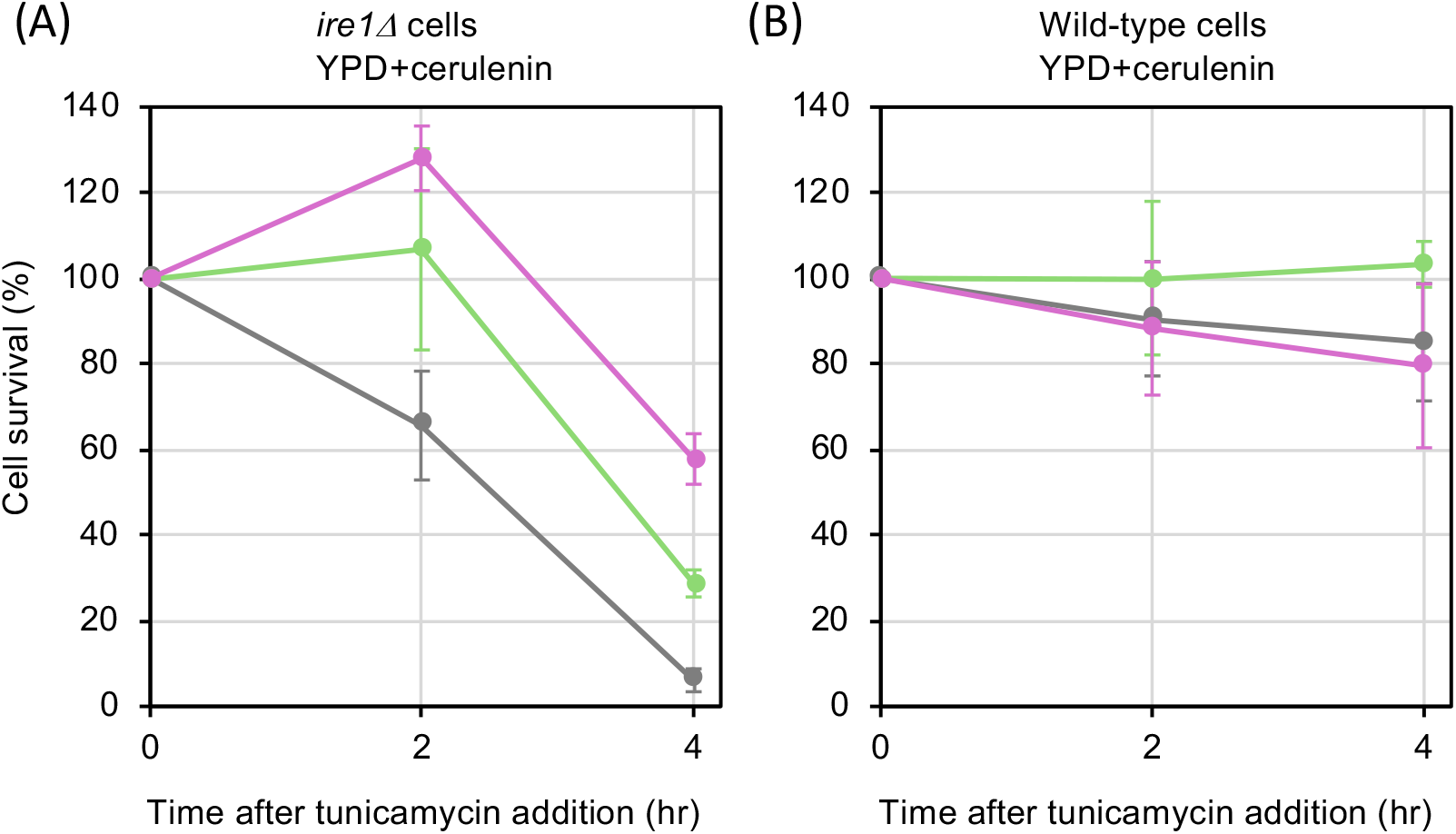
Rescue of viability of tunicamycin-treated cells by MCFs. Wild-type cells (BY4741) were cultured in fatty acid-supplied YPD containing 5.0 µg/mL cerulenin and treated with 0.6 µg/mL tunicamycin for the indicated durations. The fatty acids supplied were 500 µg/mL palmitic acid only (C16), 130 µM lauric acid with 500 µg/mL palmitic acid (C16+C12), or 130 µM capric acid with 500 µg/mL palmitic acid (C16+C10). Cell survival under the tunicamycin stress was monitored using colony formation assay.

As a possible mechanism by which MCFs mitigate ER stress, the ER membrane may exhibit altered properties in the presence of MCF-containing phospholipids. To support this hypothesis, we investigated the localization of an aberrant ER-localized transmembrane protein. The mCherry-tagged version of transmembrane methyltransferase Cho2 that carried the G102A/G104A functionless mutation (Cho2-G102A/G104A-mCherry; Ishiwata-Kimata et al., 2021) appeared to aggregate (Fig. 4A). In contrast, the MCF treatment dispersed this protein, which exhibited a typical double-ring-like ER distribution (Fig. 4B and C).

**Figure 4.**
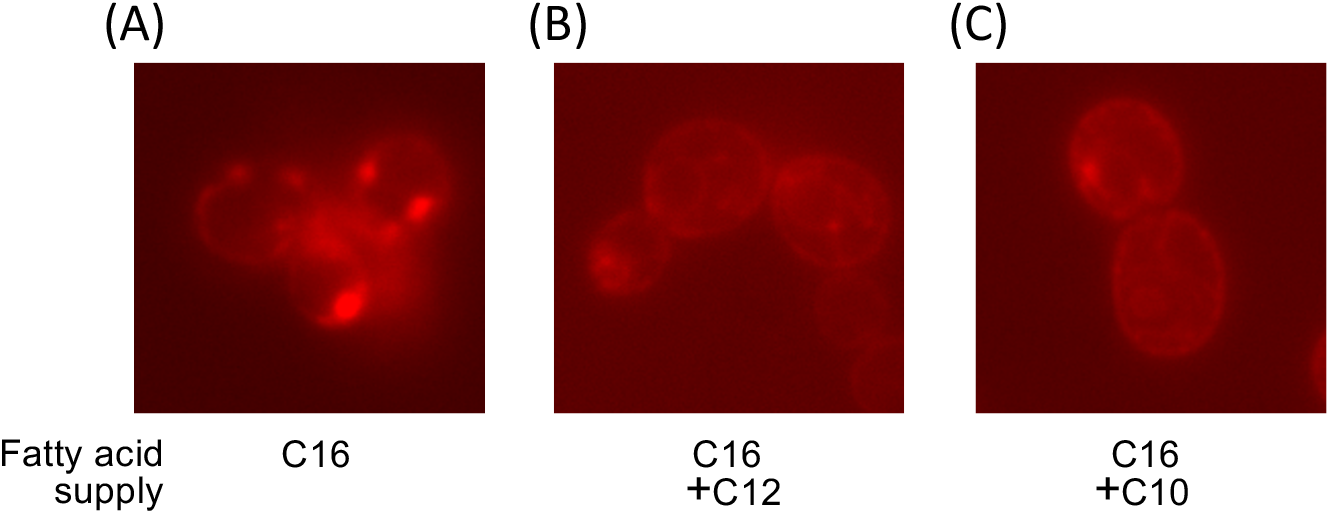
Dispersion of transmembrane protein aggregates by MCFs. Wild-type cells (BY4741) carrying pYT-CUP1p-CHO2-G102A/G104A-mCherry were cultured in fatty acid-supplied YPD containing 5.0 µg/mL cerulenin in the presence of 500 µM CuSO_4_ to induce the expression of Cho2-G102A/G104A-mCherry. The fatty acids supplied were 500 µg/mL palmitic acid only (A; C16), 130 µM lauric acid with 500 µg/mL palmitic acid (B; C16+C12), or 130 µM capric acid with 500 µg/mL palmitic acid (C; C16+C10). After 4-hr culturing, the cells were observed under a fluorescence microscope to observe m-Cherry fluorescence.

## DISCUSSION

Taken together, here we propose that, as a previously undisclosed outcome of the UPR, Hac1 induces production of MCFs, which can contribute to the mitigation of ER stress, in *S. cerevisiae* cells. We deduce that phospholipids containing short acyl chains change the properties of the ER, such as fluidity, for instance, to cope with aberrant transmembrane proteins that accumulate and aggregate in the ER. Tunicamycin causes misfolding of ER-client transmembrane proteins by inhibiting their N-glycosylation. Inositol depletion also leads to lipid membrane-related abnormality (Promlek et al., 2011), which, as shown here, mitigated by MCFs.

Phosphatidylcholine (PC) deficiency is known to induce ER stress in *S. cerevisiae* cells (Thibault et al., 2012). Bao et al. (2021) reported that PC deficiency is compensated for by genetic mutations that shorten the lipid acyl chains. A new finding in our current study is that lipids with short acyl chains are produced by the UPR, which is an intrinsic transcriptional program to cope with stress conditions.

Since, as described in the Introduction section, Hac1 controls hundreds of genes, it has not yet been determined which of them is responsible for the MCF production. Complicated changes in the metabolic network caused by UPR may result in the shortening of lipid acyl chains. Another intriguing question is whether the MCF production upon ER stress is also observed in other eukaryotic species. Further studies are required to address these issues.

## MATERIAL AND METHODS

### Yeast strains and plasmids

In all experiments shown here, we used the laboratory-standard congenic strains BY4741 (*MAT***a** *leu2Δ0 ura3Δ0 his3Δ1 met15Δ0*), BY4742 (*MAT*a *leu2Δ0 ura3Δ0 his3Δ1 lys2Δ0*), and their derivatives (Brachmann et al., 1998). The *fas2Δ::kanMX4* deletion module was PCR-amplified from the diploid *FAS2*/*fas2Δ* strain Y21061, which had been derived from EUROSCARF (http://www.euroscarf.de/), and was used for transformation of BY4741 to obtain the haploid *fas2Δ* strain BY4741-*fas2Δ. HAC1*i cells, a derivative of BY4742 carrying the *ire1Δ::kanMX4* mutation and a *HAC1* mutation to remove the intron sequence, were constructed as previously described (Nguyen PTM et al., 2022).

A DNA fragment carrying the coding sequence of a C-terminally mCherry-fused missense mutant of Pma1 (Pma1-2308-mCherry) and the *CYC1* terminator sequence was PCR-amplified from the plasmid pYT-CUP1p-PMA1-2308-mCherry (Mai et al., 2019) using the primer sets attaccggatcaattcggggATGACTGATACATCATCCTCTTCAT and ttaacaggcctgtttaaacgGCAAATTAAAGCCTTCGAGCGTCCC (the capital letters anneal to pYT-CUP1-PMA1-2308-mCherry), and was fused to the BamHI-digested YCp vector pCM189 (Garí et al., 1997) using the Gibson assembly kit (New England BioLabs). The resulting plasmid pCM189-PMA1-2308-mCherry was used to express Pma1-2308-mCherry under the control of the Tet-off promoter in *S. cerevisiae* cells (Sprengel and Hasan, 2007).

We also modified pYT-CUP1p-PMA1-2308-mCherry to replace the Pma1-2308 sequence to the Cho2-G102A/G104A-coding sequence (Ishiwata-Kimata et al., 2021). The resulting plasmid pYT-CUP1p-CHO2-G102A/G104A-mCherry was used to express Cho2-G102A/G104A under the control of the copper-inducible *CUP1* promoter in *S. cerevisiae* cells.

### Chemicals

Palmitic acid (Tokyo chemical industry), lauric acid (Tokyo chemical industry), capric acid (Tokyo chemical industry), tunicamycin (Sigma Aldrich), cerulenin (Wako), and doxycycline (Tokyo chemical industry) were prepared as stock solutions at 50 mM in ethanol, 16 mM in methanol, 16 mM in ethanol, 2 mg/ml dimethyl sulfoxide, 5mg/ml ethanol, and 1 mg/mL water, respectively, and stored at −30 °C.

### Yeast culture and stress treatments

Throughout this study, cells were aerobically cultured at 30°C in appropriate liquid media. Yeast extract/peptone/dextrose (YPD) and Synthetic Complete (SC) media were prepared as described in Adams et al. (1997). Based on the Difco manual (1984), we composed a Yeast Nitrogen Base without inositol and used it to prepare SC medium not containing inositol (SC-inositol).

BY4741 cells precultured overnight in liquid YPD were diluted to an OD_600_ of 0.4 with fresh liquid YPD and further incubated for 4 hrs in the main culture step before tunicamycin treatment and harvest. For inositol depletion, BY4741 cells precultured overnight in SC were washed with liquid SC-inositol, diluted to an OD600 of 0.4 with fresh liquid SC-inositol, and further incubated for 5 hrs in the main culture step before harvest. To express Cho2-G102A/G104A, BY4741 cells transformed with pYT-CUP1p-CHO2-G102A/G104A-mCherry were precultured overnight in liquid YPD, diluted to an OD_600_ of 0.4 with fresh liquid YPD containing 500 µM CuSO_4_, and further incubated for 4 hrs in the main culture step.

BY4741 cells transformed with pCM189-PMA1-2308-mCherry were maintained on SC agar plates containing 3.0 µg/mL doxycycline. To express Pma1-2308-mCherry, they were precultured overnight in liquid YPD (not containing doxycycline), diluted to an OD_600_ of 0.4 with fresh liquid YPD (not containing doxycycline), and further incubated for 24 hrs in the main culture step.

BY4741-*fas2Δ* cells were maintained on YPD agar plates containing 500 µM palmitic acid. They were precultured overnight in liquid YPD containing 500 µM palmitic acid, diluted to an OD_600_ of 0.4 with fresh liquid YPD containing 500 µM palmitic acid, and further incubated for 24 hrs in the main culture step before tunicamycin treatment and harvest. For the lauric acid supplementation, the culture media contained 48 µM lauric acid in the preculture step and 130 µM lauric acid in the main culture step, in addition to 500 µM palmitic acid. For the capric acid supplementation, the culture media contained 48 µM lauric acid in the preculture step and 130 µM lauric acid in the main culture step, in addition to 500 µM palmitic acid.

BY4742 and *HAC1*i cells precultured overnight in liquid SC were diluted to an OD_600_ of 0.4 by fresh liquid SC and further incubated for at least 4 hrs in the main culture step.

To treat cells with 5.0 µg/mL cerulenin, the aforementioned culturing procedures were modified as follows. In the preculture step, we used the culture media containing 500 µM palmitic acid. In the main culture step, we used the culture media containing 500 µM palmitic acid and 5 µg/mL cerulenin. For the lauric acid supplementation, the culture media further contained 48 µM lauric acid in the preculture step and 130 µM lauric acid in the main culture step. For the capric acid supplementation, the culture media further contained 48 µM lauric acid in the preculture step and 130 µM lauric acid in the main culture step.

### RNA analysis

As described previously (Le et al., 2021), total RNA was extracted from *S. cerevisiae* cells and subjected to oligo(dT)-primed reverse transcription (RT), followed by PCR using a *HAC1*-specific primer set (TACAGGGATTTCCAGAGCACG, TGAAGTGATGAAGAAATCATTCAATTC). The RT-PCR products were separated by agarose electrophoresis, and fluorescent images of the ethidium bromide-stained gels were analyzed using the ImageJ software. The *HAC1* mRNA splicing efficiency was calculated using the following formula:

100 × [band intensity of the short form/(band intensity of the long form + band intensity of the short form)].

The short form is the product from *HAC1*i mRNA, whereas the long form is the product from *HAC1*u mRNA.

### Other techniques

To monitor the lipid fatty acid composition, saponification and methylation of lipids in harvested cells were performed as described in Ishiwata-Kimata et al. (2021). Then, fatty acid methyl esters (FAMEs) were quantitively detected by gas chromatography in TechnoSuruga Laboratory Co., Ltd. (Shizuoka, Japan).

To assess cell viability, cultures were appropriately diluted in liquid YPD and spread on YPD agar plates, which were incubated at 30°C for 3 days before counting the colonies. The survival ratio in the presence of tunicamycin is expressed as the ratio of the colony formation units (CFU) at a selected time point to the CFU just before tunicamycin addition. The CFU was corrected against the OD_600_ of the culture because the cells continued to grow even in the presence of tunicamycin.

Fluorescence microscopy to observe mCherry fluorescence was performed as described in Mai et al. (2019).

### Statistical analysis

Data from three independent determinations are expressed as the average and standard deviations.

